# Phosphoproteomics Reveals AMPK Substrate Network in Response to DNA Damage and Histone Acetylation

**DOI:** 10.1101/2020.07.02.182121

**Authors:** Yuejing Jiang, Xiaoji Cong, Shangwen Jiang, Ying Dong, Lei Zhao, Yi Zang, Minjia Tan, Jia Li

## Abstract

AMPK is a conservative energy sensor that plays roles in diverse biologic processes via direct phosphorylation on various substrates. Emerging studies have demonstrated the regulatory roles of AMPK in DNA repair, but the underlying mechanisms remain to be fully understood. Herein, using mass spectrometry-based proteomic technologies, we systematically investigate the regulatory network of AMPK in DNA damage response. Our system-wide phosphoproteome study uncovers a variety of newly-identified potential substrates involved in diverse biologic processes, whereas our system-wide histone modification analysis reveals a linkage between AMPK and histone acetylation. Together with these findings, we discover that AMPK promotes apoptosis by phosphorylating ASPP2 in irradiation-dependent way and regulates histone acetylation by phosphorylating HDAC9 in irradiation-independent way. Besides, we reveal that disturbing the histone acetylation by the bromodomain BRD4 inhibitor JQ-1 enhanced the sensitivity of AMPK-deficient cells to irradiation. Therefore, our studies provided a source to study the phosphorylation and histone acetylation underlying the regulatory network of AMPK, which could be beneficial to understand the exact role of AMPK in DNA damage response.

## Introduction

5’AMP-activated kinase (AMPK) is a heterotrimeric serine/threonine kinase complex with one catalytic subunit α (including α1 and α2) and two regulatory subunits β (including β1 and β2) and γ (including γ1, γ2 and γ3), which contains theoretically 12 different types of heterotrimers. As a central metabolic sensor to restore intercellular energy homeostasis, activation of AMPK switches off anabolic pathways that consume ATP and switches on catabolic pathways that generate ATP. There are two coordinated mechanisms that modulate the activity of AMPK. One is phosphorylation at its conserved Thr172 in the activation loop of AMPKα1 subunit, which is regulated by upstream kinases LKB1 and CaMKKβ. The other is the binding of ADP/AMP to γ subunit to allosterically activate AMPK by stabilizing it in an active conformation [1]. Once activated, AMPK, in turn, further phosphorylates diverse downstream substrates, such as metabolic enzymes, transcription factors and co-activators, to balance energy homeostasis either by short-term provocations of metabolic signaling cascades or by long-term regulations of transcription and posttranslational modification [2].

Emerging studies have uncovered the contradictory roles of AMPK in tumor development and cancer therapy. AMPK was reported to be a key mediator contributing to the suppression effect of LKB1 signaling cascades [3, 4]. Such hypothesis is supported by a series of studies from metformin, whose mechanism of action is widely-recognized to be associated with AMPK activation [5, 6]. Metformin could significantly inhibit tumor growth and improve chemo-sensitivity and radio-sensitivity via AMPK activation in various types of tumor [7-10]. Remarkably, several studies showed the effect of metformin to kill the cancer stem like cells [11-13], which are considered a major barrier in cancer therapy. Studies from another AMPK activator AICAR [14] also supported such hypothesis. One the other hand, other studies suggest AMPK could be a context-dependent tumor promoter. The biological consequences in which AMPK is supra-physiologically activated by compounds are different from those in which AMPK is physiologically activated by cellular stresses [15]. Indeed, AMPK energy-sensing pathway supports cells to survive in hypoxic and nutrient-deficient conditions, which is regarded as a typical tumor microenvironment. Supporting such assumption is that autophagy, which is partially considered to be regulated by AMPK, might provide enough nutrients to support cancer survival by degrading cellular organelles or proteins [16]. In addition, the GAPDH generated by fatty acid oxidation, which is upregulated by AMPK activation, is beneficial to protect cancer cells from oxidative stresses by neutralizing cytotoxic ROS [17]. Furthermore, increasing phospho-AMPK levels is associated with higher tumor grade in prostate cancer [18]. Therefore, these contradictory findings suggest an urgent need to understand the exact role of AMPK in tumor biology.

The above puzzled phenomenon could be partially explained by the complicated functional consequences caused by the diverse substrates of AMPK. For example, AMPK-mediated phosphorylation on p53 [19-21] and mTORC1 [22-24] signaling pathway is essential for activator-induced tumor suppression. In contrast, AMPK could promote tumor growth via other mechanisms, including the induction of mitophagy by phosphorylating ULK1 [16], the upregulation of stress-induced gene transcription by phosphorylating histone H2B [25] and the promotion of FA oxidation to neutralize oxidative stress [17]. Moreover, new substrates identified in mitochondrial fission [26, 27], hippo-YAP signaling pathway [28] and mitosis [22] implied a subtle regulatory role of AMPK in cancer. In addition, the pharmacological effects of two anti-cancer drugs, etoposide and cisplatin, were reported to be partially dependent on AMPK activation [29-31]. Despite these understanding, detailed mechanism is not yet fully understood how AMPK coordinately regulated these cascades and whether some unknown partners were involved in this process. Indeed, our knowledge of the cancer-associated AMPK substrates is still limited. Therefore, a comprehensive understanding of the substrates of AMPK is a critical step toward understanding the role of AMPK in cancer.

Human tumors share various biological hallmarks acquired during the multistep development. One of these hallmarks is genome instability, which is acquired during cancer development and drug resistance [32]. Abnormal or deficient DDR results in genomic instability and neoplastic transformation. In recent years, emerging reports suggested that AMPK was involved in DDR but the regulatory role remained to be fully understood [33-35].

To address this question, we carried out a system-wide phosphoproteome study by mass spectrometry-based proteomic technologies. Our results showed that a variety of newly-identified substrates were involved in diverse biologic activities, which shed light on a broad and complex regulatory network of AMPK in DDR. In addition, our system-wide histone modification analysis showed that AMPK played a role in modulating global histone acetylation levels. Disturbing the histone acetylation by the bromodomain BRD4 inhibitor JQ-1 enhanced the sensitivity of AMPK-deficient cells to irradiation via induction of apoptosis. Thus, our studies provided an abundant source to study the phosphorylation and histone acetylation underlying the regulatory network of AMPK, which might be beneficial to understand the exact role of AMPK in tumor biology.

## Results

### Establishment of AMPKα1/α2-double knockout cell line by TALENs

We first investigated whether AMPK was involved in DDR. X-ray irradiation induced cellular DNA damages and activated DDR signaling to execute DNA double strand breaks (DSBs) repair. After single dose of X-ray irradiation exposure, both AMPK activation signal phosphor-AMPK (Thr172) and its substrate signal phosphor-ACC (Thr79) increased as irradiation dose elevated, suggesting that AMPK activated in a dose-dependent way in DDR (Figure 1A and B). To investigate the underlying mechanisms of AMPK activation during DDR, we established stable AMPKα1/α2 (two AMPK catalytic isoform)-double knockout mouse embryonic fibroblasts (MEFs) cell line using TALENs (Transcription Activator-Like Effector Nucleases) technology. TALENs technology is a genome-editing method widely used to generate knockout *C. elegans*, rats, mice and zebrafishes [36-42]. It is also used in genomic modification of human embryonic stem cells and induced pluripotent stem cells (IPSCs). The establishment method was according to previous reports [39, 41, 43]. We obtained 3 stable AMPKα-KO cell lines (named as 1#, 2# and 3#). The AMPKα1 and α2 protein expression levels could not be detected by immunoblot, suggesting successful deletion of total AMPKα subunits (Figure 1C). To evaluate whether downstream signaling was also impaired, we treated cells with AMPK activator AICAR. In contrast to wildtype MEFs cells, AICAR treatment didn’t induce AMPK activation signals phospho-AMPKα (Thr172) and substrate signals phospho-ACC (Thr79) in 3 knockout cell lines, further suggesting the loss of AMPKα1 and α2 kinase activity (Figure 1C). Besides, we extracted genomic DNA from individual cell lines for sequencing and found nucleotide deletion in leading chain and lagging chain. In addition, our genomic sequencing of these cell lines also showed nucleotide deletion happened in both leading chain and lagging chain in all 3 cell lines (Figure S1). In contrast to 3# cell line, TALENs induced non-3 nucleotides-deletion in 1# and 2# cell line that resulted in absolute target knockout in genomic level because of coding frame shift. We next chose 1# and 2# cell lines for further functional assay. To address the issue whether AMPK is involved in DDR, we exposed wildtype MEF, 1# and 2# cells to a single dose of ion irradiation. We observed an increasing γH2A.X (phosphor-H2A.X Ser139) foci (a marker of DSB measured by immunofluorescence) after irradiation exposure (Figure 1D and E). Comparable percentage of γH2AX foci-positive cells at 48 hours post-irradiation in 1# and 2# cell line suggested these two cell lines could be functionally equivalent in DDR. Therefore, 1# cell line (defined as AMPKα-KO cells in this paper) was selected for further proteome analysis. Meanwhile, compared with wildtype MEFs, more γH2AX foci remained at 48 hours in 1# and 2# cell lines, suggesting AMPKα was required to promote DSBs repair efficiency. Besides, a prolonged G2 phase arrest was observed in AMPKα-KO cells after irradiation (WT, 0 hr 27.30% to 24 hr 25.72%, p =0.690; KO, 0 hr 30.55% to 24 hr 40.65%, p =0.011) (Figure 1F and G). Taken together, these results suggested the kinase activity of AMPKα was required in DDR, but the underlying regulatory network remained to be understood.

**Figure 1.**
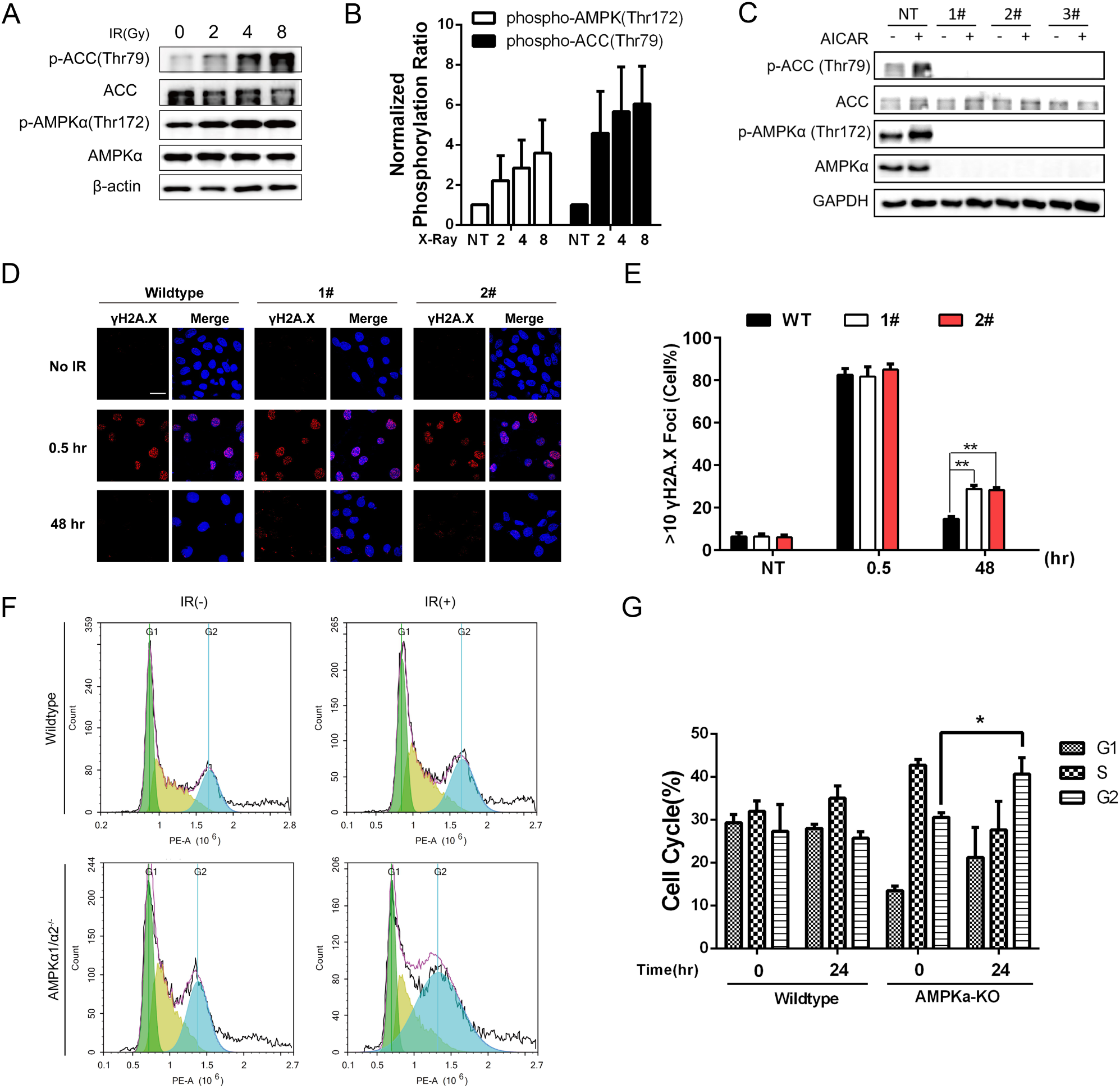
AMPK plays biologic roles in DNA damage response. **A**. AMPK activated in a dose-dependent manner. The wildtype MEFs cells were exposed to increasing does of X-ray (0, 2, 4, 8 Gy) and sampled at 10 mins after irradiation. **B**. Quantitative result of panel A as measured by ImageJ software. Mean values ±SEM. from three independent experiments are shown. **C**. Validation of AMPKα1/α2-KO cell lines 1#, 2# and 3#. Parental wildtype MEFs cells and three AMPKα1/α2 double knockout cell lines were treated with or without 1 mM AICAR for 30 mins. AICAR, 5-aminoimidazole-4-carboxamide-1-β-D-ribofuranoside, AMPK activator. **D**. AMPK knockout impairs efficient repair. The wildtype and AMPKα1/α2-KO MEFs cells 1# and 2# were exposed to a single dose of 4 Gy X-Ray. After recovery for 0.5 or 48 hours, the cells were fixed and stained with anti-γH2A.X antibody and observed under microscopy. Scale bar, 10 μm. **E**. Quantitative results measured in panel A. Cells with more than ten γH2A.X. foci were regarded as positive ones. More than 100 cells in each group were imaged and counted. Mean values ±SEM. from two independent experiments are shown. **F**. AMPK knockout prolonged G2 phase arrest. The wildtype and AMPKα-KO MEFs cells were treated with or without 10 Gy X-Ray and cultured for 24 hours before assay. **G**. Quantitative results measured in panel F. Mean values ±SEM. from three independent experiments.

### Quantitative phosphoproteomics uncovered broad signaling pathways regulated by AMPK in DNA damage response

To further profile the regulatory networks of AMPK in DDR, we carried out a system-wide phosphoproteome analysis (Figure 2A). Previous studies suggested that more abundant signaling pathways would be activated in response to irradiation at relatively high dose than low dose [44, 45]. According to the result that AMPK activated and phosphorylated substrates in a dose-dependent manner (Figure 1A and B), we hypothesized AMPK likely activated more signaling pathways at higher irradiation dose. Therefore, a relatively high dose (10 Gy) that mimicked the stressed cellular microenvironment in radio-combined cancer therapy was selected in further experiment. The proteome of AMPKα-KO cells was labeled with “heavy” (^13^C_6_-Lys and ^13^C_6_^15^N_4_-Arg) amino acids, whereas the proteome of the wild type cells was labeled with “light” (^12^C_6_-Lys and ^12^C_6_^13^N_4_-Arg) amino acids in cell culture. AMPKα-wildtype and AMPKα-KO cells were prepared in both basal group and irradiation group at the same time. After irradiation, cells from irradiation group were released for 90 minutes before harvest to fully activate diverse signaling pathways involved in DNA damages response. Proteins were extracted from the two cell populations and mixed equally for further analysis. Two types of protease, trypsin and chymotrypsin, were used to improve the sequence coverage of the phosphoproteome as previously reported [46]. Collectively, we identified 11,065 phosphosites with a localization probability higher than 0.75 in the basal and irradiation group. First, we verify the reproducibility between replicate experiments in basal (trypsin and chymotrypsin digested) and irradiation (trypsin and chymotrypsin digested) group (Figure S2). Scatter plot depicting the correlation between two replicates in each group showed high correlation coefficient value (r = 0.78-0.84), suggesting reliable reproducibility of our phosphoproteomics data. We used a criterion of more than 1.5-fold between basal and irradiation groups to define those significantly different phosphosites (p-value <0.05 by student’s t-test). 539 down-regulated phosphosites and 389 up-regulated phosphosites were identified in the basal group (Figure 2B and Table S1). Phosphosites of well-known AMPK substrates was found significantly downregulated in AMPKα-KO cells (Figure 2B). Motif analysis showed that consensus AMPK substrate motif could be enriched in the downregulated phosphosites in AMPKα-KO cells (Figure S3). Together, these results suggested the high-quality of our proteomics data.

**Figure 2.**
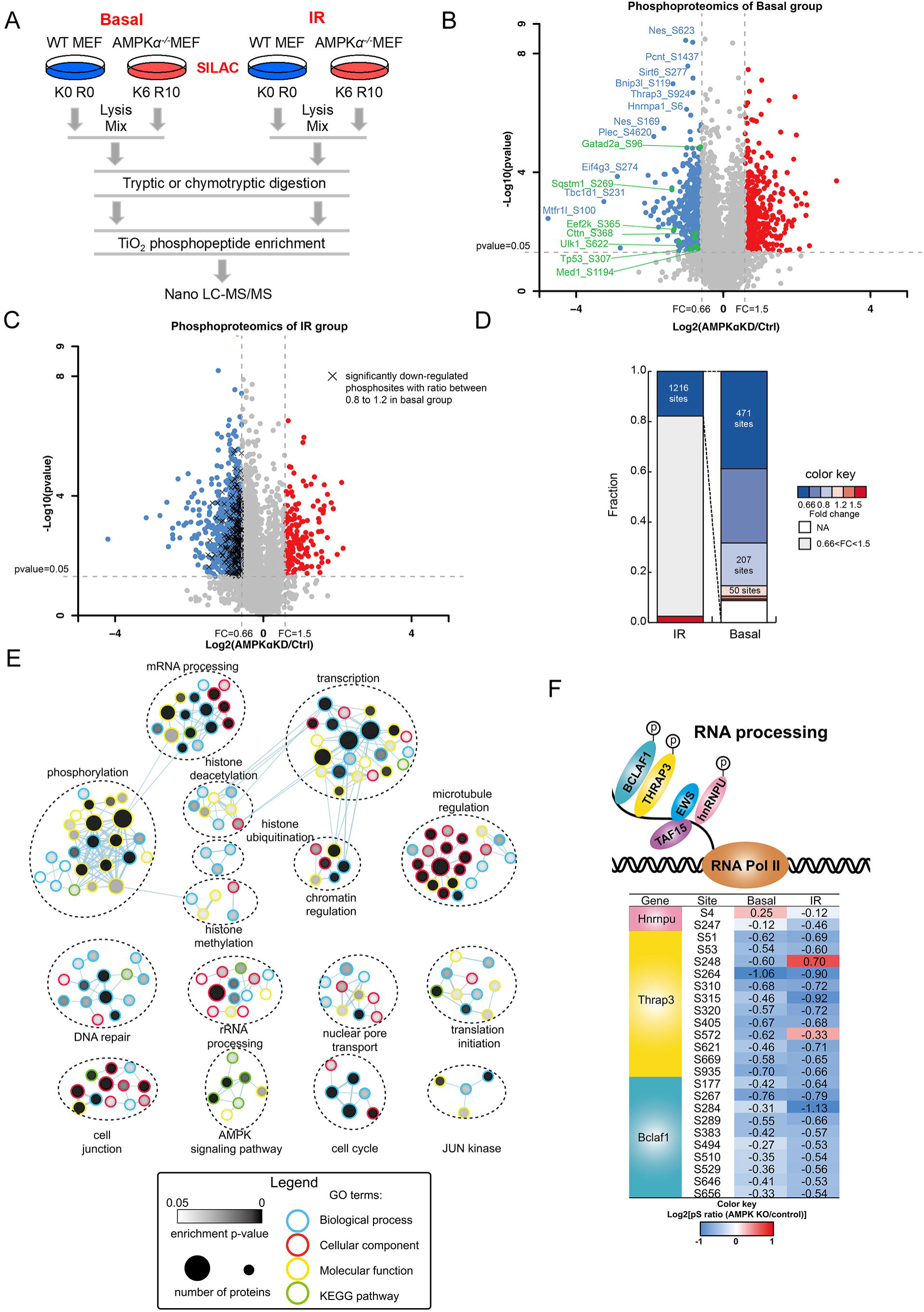
Quantitative phosphoproteomics uncovered broad signaling pathway regulated by AMPK in DNA damage response. **A**. General workflow of quantitative phosphoproteomic analysis. The SILAC-labeled wildtype and AMPKα-KO MEFs cells in basal or IR group were lysed and mixed together. Phosphopeptide were enriched and analyzed by Nano LC-MS/MS. Two replicates were analyzed. **B**. Volcano plot of quantitative phosphoproteomics analysis of basal group. The phospho-peptides with fold change (AMPKα-KO/wildtype) >1.5 (p-value < 0.05) were selected as significant regulated. Red dots were the phosphosites significantly up-regulated. Blue dots are the phosphosites significantly down-regulated. Green dots were the phosphosites on known AMPK substrates. **C**. The bar graph shows changes of upregulated phosphorylation sites after ion irradiation in basal group. **D**. Volcano plot of quantitative phosphoproteomics analysis of IR group. Cross represented significantly down-regulated phosphosites in IR group while these phosphosites were unchanged in basal group (0.8<FC<1.2). **E**. Representative enrichment results from phosphoproteomic analysis. Phosphorylation significantly downregulated proteins after ion irradiation were analyzed for GO term and KEGG pathways enrichment using DAVID. Visualization of results was performed with Cytoscape and EnrichmentMapApp. Nodes represent a gene set with enriched GO terms or KEGG pathways. Edges represent sharing proteins among nodes. Terms visualized have a p-value cutoff of < 0.05. **F**. The diagram shows significantly changed phosphosites in RNA processing.

Meanwhile, 1,216 down-regulated phosphosites and 169 up-regulated phosphosites were identified in the irradiation group (Figure 2C-D and Table S1). This result indicated various protein phosphorylation events could be regulated by AMPK during DDR. Of these significantly downregulated phosphosites, 471 phosphosites (38%) were also downregulated (FC <0.667) in basal group (Figure 2C). In contrast, 257 phosphosites (21%) remained unchanged (0.8< FC <1.2) in basal group (Figure 2C and D). For example, phosphorylation on Xrcc1 Thr452 was slightly downregulated in basal group (log2FC =−0.28, p-value =0.01) while it was significantly altered after irradiation (log2FC =−0.78, p-value =0.005).Therefore, these results suggested that AMPK played additional regulatory roles when cells suffered severe genomic stress, as compared to its routine regulatory roles in basal status.

To gain insight into the possible biological functions of AMPK during DDR, we subjected the all significantly down-regulated and up-regulated phosphoproteins identified in irradiation group to bioinformatic enrichment analysis using GO and KEGG databases by the DAVID bioinformatic tool. In consistent with a prolonged G2 phase arrest observed in AMPKα-KO cells after irradiation (Figure 1F and G), we discovered the changed phosphoproteins were significantly enriched in DNA damage repair-associated events, such as cell cycle, DNA repair and transcription (Figure 2E). Surprisingly, chromatin-associated functions were also highly enriched, such as chromatin regulation and histone modification (including deacetylation, ubiquitination and methylation), suggesting a potential regulatory role of AMPK in epigenetic modification. Meanwhile, we also discovered a potential crosstalk between DNA damage and RNA processing (both mRNA processing and rRNA processing), in consistent with emerging evidences that mRNA processing factors involved in DNA damage signal transduction. Theoretically, BCLAF1 assembled at damage sites via interacting with core spliceosome components while THRAP3 was excluded from damage sites as a consequence of transcription repression [47]. Phosphorylation events was one of factors to regulate the assemble and dissemble process. Significantly upregulated phosphosites Ser248 and Ser572 on THRAP3, as well as significantly downregulated phosphosite Ser284 on BCLAF1 were observed. Besides, mRNA splicing factor hnRNPUL1, a DNA-end resection regulator, was found significantly downregulated at Ser4 during DDR (Figure 2F). These findings implied the phosphorylation events on these mRNA splicing factors might be involved in DSB repair. Taken together, bioinformatic enrichment analysis revealed a broad and complicated regulatory role of AMPK involved in diverse signaling pathways.

### Bioinformatic analysis identifies ASPP2 as a new substrate of AMPK involved in apoptosis

We next analyzed the significantly changed phosphoproteins in irradiation group by Motif-X algorithm to predict the preferred motif sequences. It was well established that AMPK phosphorylated diverse substrates at consensus amino acid sequence motif. Briefly, −5 site and +4 site contained a hydrophobic amino acid like L/M/V/I/F (Φ), while −3 or −4 site contained at least one basic amino acid R/K (β) [48]. Motif enrichment analysis of downregulated phosphosites in IR group was carried out. Among several enriched motifs proposed by Motif-X algorithm (Figure 3A), the sequences ranked 2^nd^ and 3^rd^, xxxxRRxSxxxxxxx and xxxxRxxSxxxxxxx, partially matched to the classical AMPK substrate motif ΦxβxxSxxxΦ. This possibly indicated that there existed some potential AMPK substrates not strictly match classical motif very well.

**Figure 3.**
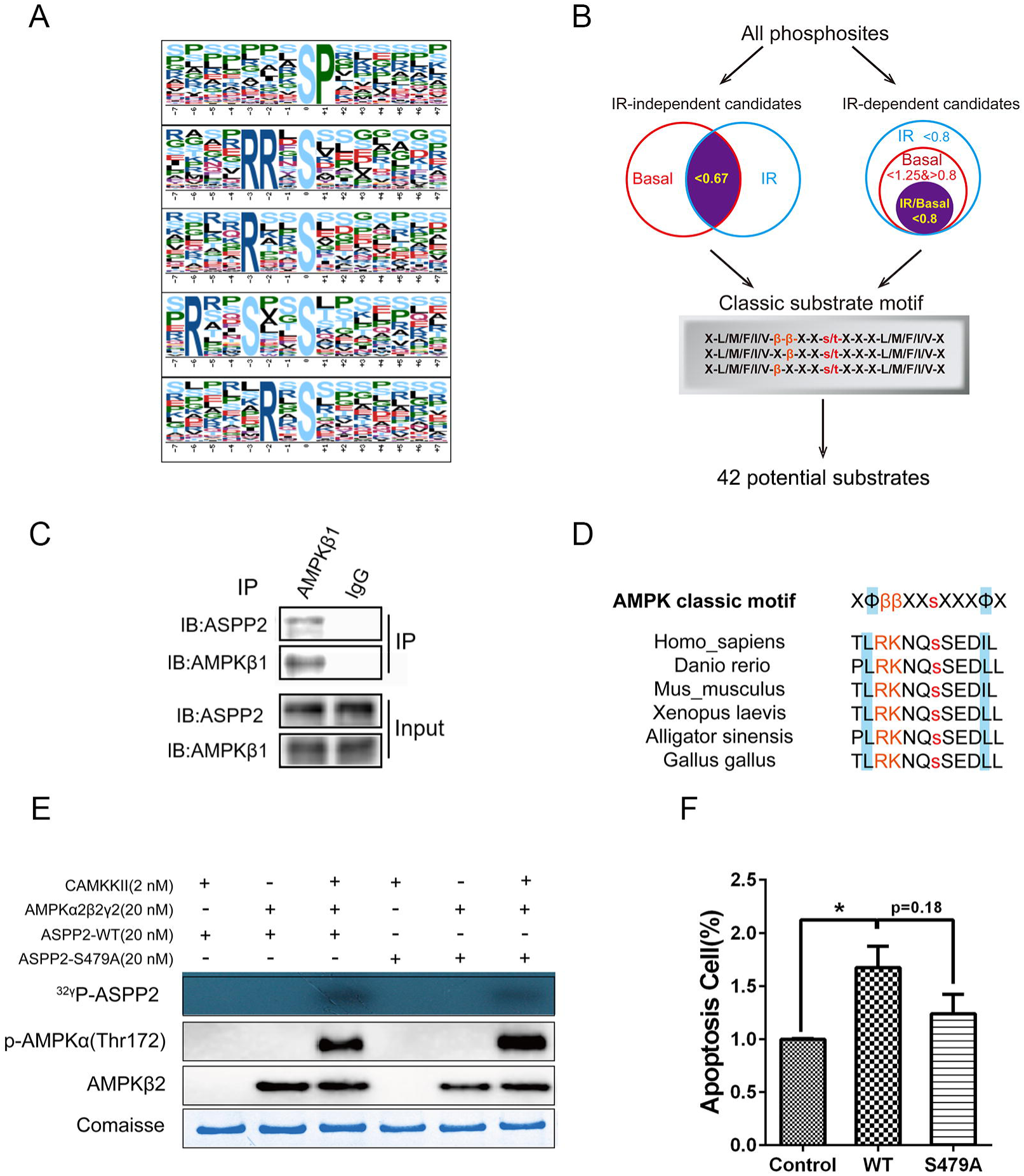
Bioinformatic analysis identified ASPP2 as a novel substrate of AMPK involved in apoptosis. **A**. Motif analysis of significantly downregulated in IR group. Motif-X was used to analyze the phosphorylation motif. **B**. Strategy diagram for candidate AMPK substrate identification. **C**. Endogenous co-immunoprecipitation using anti-AMPKβ1 antibody discovered interaction of ASPP2 and AMPKβ1 in 293T cells. **D**. The evolutionally conserved amino acid sequence surrounding Ser479 on ASPP2 matches the consensus AMPK phosphorylation motif. **E**. AMPK phosphorylates ASPP2 at Ser479 in vitro. In vitro kinase assays were performed using 200 nM ASPP2-(1-765 aa) truncation, 20 nM AMPKα2β2γ2 (inactive or active), 0.1 mM DTT and 5 μCi γ-[^32^P] ATP per reaction. **F**. Ser479 phosphorylation is required in ASPP2-induced apoptosis. HeLa cells were transfected with plasmids and etopic overexpressing wildtype ASPP2-WT or phosphonull ASPP2-S479A mutants for 48 hours. Next, the cells were harvested and stained by PI and FITC-AnnexinV dyes and the fluorescence intensity was examined by flow cytometer. Data are presented as a densitometric ratio change that normalized to the negative control.

Among the above significantly downregulated phosphosites, we further divided them into two groups, IR-independent substrates and IR dependent substrates. The IR-dependent substrates (DNA-damage-associated substrates) were phosphorylated by AMPK under DDR. These phosphosites were downregulated in irradiation group but remained nearly unchanged in basal group (normalized heavy/light ratio, 0< IR/Basal <0.8 and 0.8< Basal <1.25). In contrast, the IR-independent substrates (general substrate candidates) were phosphorylated by AMPK independent of DDR. These phosphosites were significantly downregulated in either basal group or irradiation group (heavy/light ratio <0.67). To identify direct potential substrates of AMPK, we aligned these phosphosites in both IR-dependent and IR-independent substrates to AMPK consensus motif. Based upon the criteria, we totally obtained 42 potential substrates (Figure 3B and Table 1).

**Table 1.**
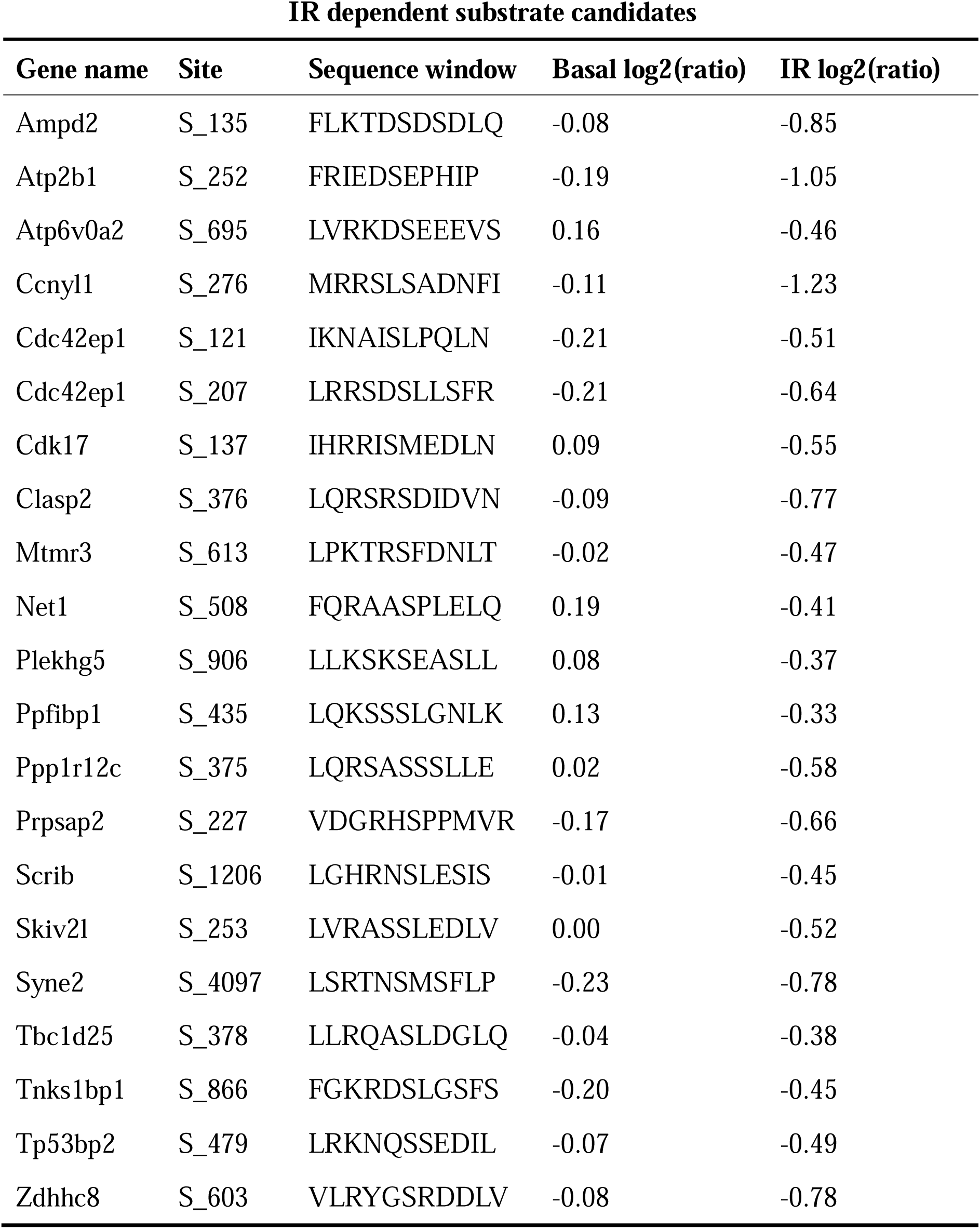
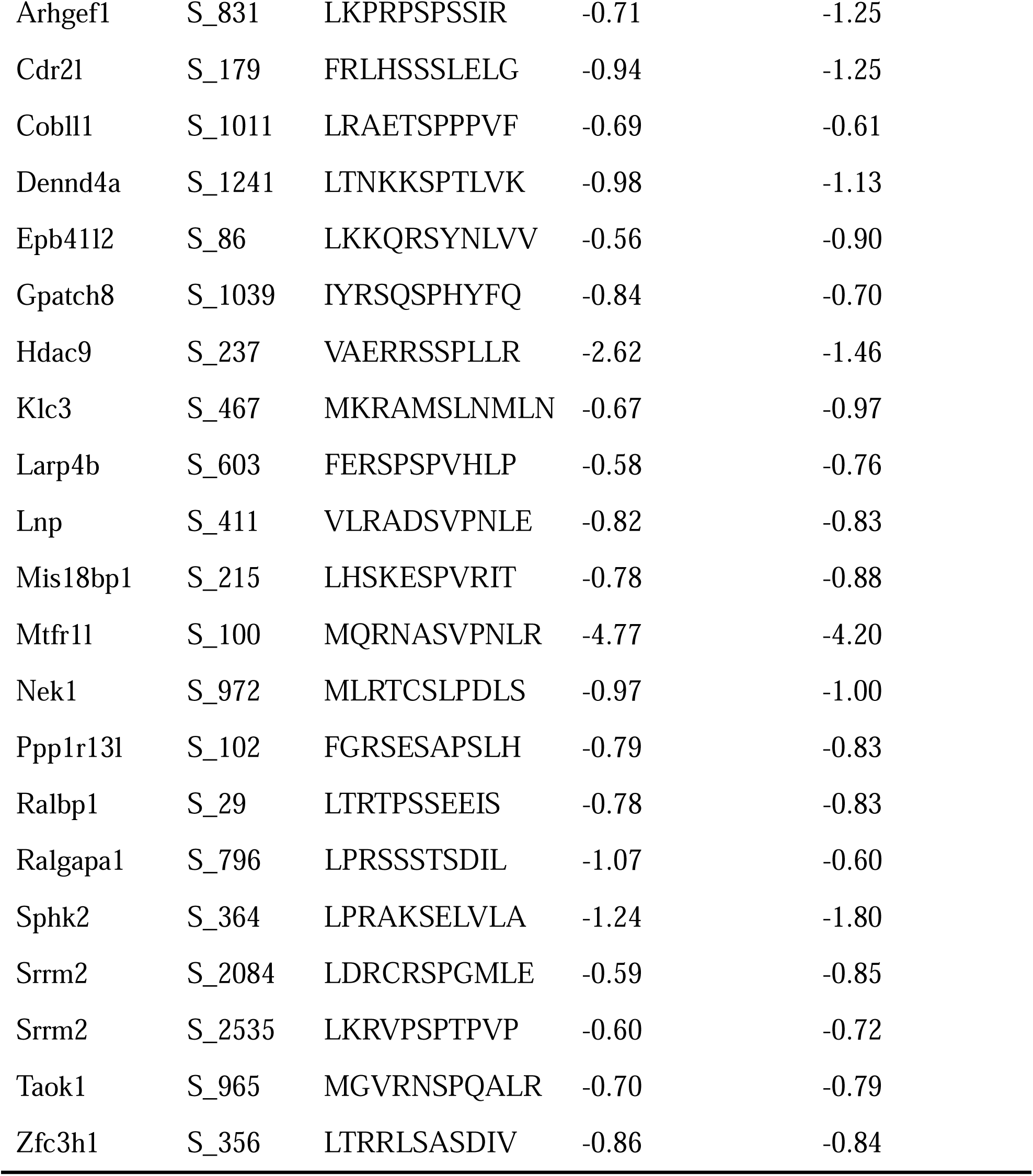
The candidate AMPK substrates identified in quantitative phosphosproteomics

To further confirm the reliability of our results, we randomly selected one of the substrates, apoptosis-stimulating of p53 protein 2 (ASPP2) for further validation. Ser479 on ASPP2 was considered as DNA-damage-associated substrate because the normalized IR/basal ratio of Ser479 on ASPP2 was 0.78 (normalized heavy/light ratio, Basal =0.95, IR =0.74). p53 was a well-established tumor suppressor involved in both cell cycle arrest and apoptosis. ASPP2 was initially identified as a binding partner of p53 and required in p53-mediated apoptosis [49, 50]. We first investigated whether protein interaction existed between AMPK and ASPP2. Plasmids expressing Flag-ASPP2 and Myc-AMPKβ1 were synchronically introduced into 293T cells for 48 hours and the whole cell lysates were subjected to exogenous immunoprecipitation with the anti-Flag antibody. Immunoblot showed interaction between AMPKβ1 and ASPP2 (Figure S4). The endogenous protein-protein interaction was further confirmed in 293T cells in immunoprecipitation assay using AMPKβ1 antibody (Figure 3C). Bioinformatic analysis revealed that amino acid sequence surrounding Ser479 on ASPP2 matched the AMPK substrate motif and kept conservative during species evolution (Figure 3D). Therefore, we investigated whether AMPK could phosphorylate ASPP2 at Ser479 by *in vitro* kinase assay using [^32^P]-labelled ATP. Wildtype ASPP2-(1-765 aa) truncation and mutant ASPP2-(1-765 aa) S479A (Serine converted to non-phosphorylatable alanine) that mimicked phosopho-null status were purified from *E. coil* and incubated with AMPKα2β2γ2 complex in kinase reaction buffer for 1 hour. AMPKα2β2γ2 complex with no kinase activity was purified from *E. coil*. but retained catalytic activity after incubation with upstream kinase CAMKKβ. Robust [^32^P] signal was induced by activated-AMPKα2β2γ2 complex on wildtype ASPP2 truncation (1-765 aa) but strongly attenuated on S479A mutant, suggesting AMPK phosphorylated ASPP2 dominantly at Ser479 (Figure 3E). However, weak [^32^P] signal could be detected on mutant ASPP2-S479A also implied other unknown AMPK phosphorylation sites on ASPP2 remains to be further investigated. We then investigated whether Ser479 phosphorylation was involved in apoptosis. HeLa cells were transfected with full-length wildtype ASPP2 or mutant ASPP2-S479A plasmids for 24 hours. Apoptosis cells was labeled with Annexin V or PI dyes and measured by flow cytometry. As normalized to negative control, overexpression of wildtype ASPP2 induced a 1.67-fold increased apoptosis. Compared to wildtype, overexpression of mutant ASPP2-S479A moderately impaired apoptosis (Figure 3F). Taken together, the data suggested that AMPK-mediated phosphorylation at Ser479 on ASPP2 was required in stimulation of cell apoptosis.

### Histone modification analysis characterized AMPK as a regulator of histone acetylation

Since chromatin-associated functions were highly enriched in bioinformatic analysis, we hypothesized that AMPK was involved in histone posttranslational modifications and epigenetic regulation. To address this question, we carried out system-wide analysis to investigate the histone modification by mass spectrometry in both non-irradiation basal status and irradiation stress status (Figure 4A). The heavy-labeled AMPKα-KO cells and light-labeled wildtype cells were mixed at 1:1 ratio and core histones (Histone H2A, H2B, H3 and H4) were extracted and separated with SDS-PAGE. Histones were in-gel digested with trypsin into peptides and then subjected to mass spectrometry analysis. The modification peptides were checked manually and quantified based on peak area integral according to the previous report [51]. Interestingly, AMPKα knockout resulted in significant upregulation of global histone acetylation levels (Figure 4B and Table S2). Immunoblot assay also suggested increased acetylation level at H2BK12, H3K18 and H4K16 in 1# AMPKα-KO cells, as well as in 2# AMPKα1/α2 double knockout cell line (Figure 4C). These data suggested AMPK played a negative regulatory role in global histone acetylation.

**Figure 4.**
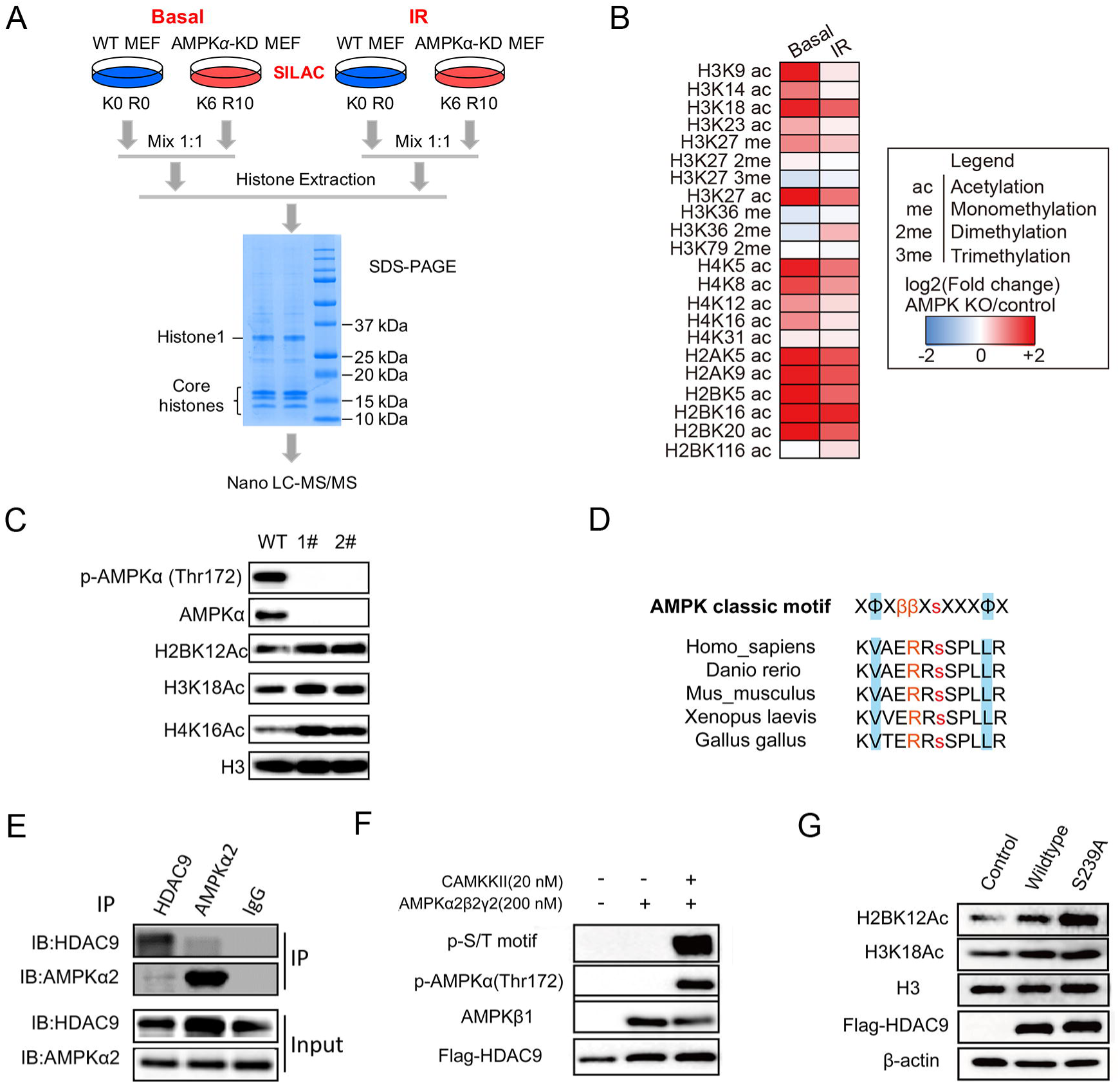
Histone modification analysis characterized AMPK as a regulator of histone acetylation. **A**. General workflow for quantification of histone modification. The SILAC-labeled wildtype and AMPKα-KO MEFs cells in basal or IR group were lysed and mixed together. Histones were extracted and analyzed by Nano LC-MS/MS. **B**. The quantified histone marks in response to DNA damage in AMPKα-KO MEFs cells. The fold change of histone posttranslational modifications in wildtype compared with AMPKα-KO MEFs cells in basal or IR group was detected by LC-MS/MS. **C**. AMPK knockdown promoted histone acetylation. Both wildtype and AMPKα1/α2 double-knockout cells (1# and 2# cell lines) were subjected to immunoblot, to validate the results presented in panel B. **D**. The evolutionally conserved amino acid sequence surrounding Ser239 on HDAC9 matches the consensus AMPK phosphorylation motif. **E**. Endogenous co-immunoprecipitation using anti-AMPKα2 and anti-HDAC9 antibodies discovered reciprocal interaction of ASPP2 and AMPKβ1 in 293T cells. **F**. AMPK phosphorylates HDAC9 at Ser239 in vitro. In vitro kinase assays were performed using immunoprecipitated Flag-HDAC9, 20 nM AMPKα2β2γ2 (inactive or active), 0.1 mM DTT, and 0.1 mM ATP per reaction. The reaction sample was subjected to immunoblot, and the signal was determined by pan phosphor-AMPK substrate antibody. **G**. Phosphonull Ser239 mutant affected the H2BK12 acetylation. 293T cells were transfected with plasmids and cultured for another 48 hours. The samples were examined by immunoblot.

To find out underlying regulation mechanism, we chose another potential substrate HDAC9 from our phosphoproteome data for investigation. The amino acid sequence surrounding Ser239 of HDAC9 matched the AMPK substrate motif and is evolved conservatively (Figure 4D). Plasmids expressing Flag-HDAC9 and Myc-AMPKα2 were synchronically introduced into 293T cells for 48 hours and the whole cell lysates were subjected to exogenous immunoprecipitation assay using the anti-Flag antibody. AMPKα2 was detected using anti-Myc antibody in Flag-HDAC9 immunoprecipitated complex suggested AMPKα2 interacted with HDAC9 (Figure S5). Endogenous immunoprecipitation assay conducted in 293T cells discovered a reciprocal interaction between AMPKα2 and HDAC9 cells (Figure 4E). Additionally, the exogenous fusion protein Flag-HDAC9 was enriched and extracted by anti-Flag antibody from 293T cells and incubated with activated or non-activated AMPKα2β2γ2 complex in kinase reaction buffer for 1 hour. Phosphorylation signal (as measured by pan phosphor-AMPK-substrate antibody) was detected on HDAC9 when incubated with activated-AMPKα2β2γ2 complex in the presence of CAMKKβ (Figure 4F). Taken together, HDAC9 was newly identified as substrate of AMPK. Afterwards, two plasmids expressing wildtype HDAC9 and phospho-null mutant HDAC9-S239A were constructed and introduced into 293T cells. Compared with negative control, overexpression of wildtype HDAC9 enhanced H2BK12 acetylation and H3K18 acetylation. In contrast, overexpression of phospho-null mutant HDAC9-S239A enhanced higher H2BK12 acetylation level but had no impact on H3K18 acetylation level (Figure 4G). Therefore, these results suggested that HDAC9 was involved in balance of histone acetylation while Ser239 phosphorylation on HDAC9 played a negative role in regulation of H2BK12 acetylation. Since there was no significant difference in HDAC9-Ser239 phosphorylation between basal and irradiation groups, AMPK likely phosphorylated HDAC9 at Ser239 in irradiation-independent manner.

Taken together, quantitative phosphosproteome and histone modifications analysis uncovered subtle crosstalk between AMPK-mediated phospho-regulation and histone acetylation.

### JQ-1 synergizing with AMPK inhibitor induces cell apoptosis in irradiation

Histone acetylation had impact on localized chromatin structures and played important roles in initiation and transduction of DNA damage signaling. Brd4, a member of the bromodomain and extraterminal (BET) family that binds acetylated histones H3 and H4, and regulates gene expression [52-54]. In addition to transcriptional regulation, BRD4 is essential in DDR and mediates the recruitment of chromatin-based repair proteins. Once sensing genome-wide DNA breaks, enhanced H4 acetylation led to Brd4 accumulation at breaks [55, 56]. Since increased acetylation level at H3K14Ac and global H4 acetylation level was observed in AMPKα-KO cells in basal group, but the tendency slightly attenuated after irradiation (Figure 4B), we hypothesized a potential linkage between AMPK and Brd4 during DDR. JQ-1, a BET inhibitor that impeded BRD4-acetylated lysine interaction, displayed anti-tumor activity in various types of cancer [57, 58]. Interestingly, JQ-1 treatment made AMPKα-KO cells more sensitive to irradiation than wildtype cells (Figure 5A). Similarly, we found JQ-1 treatment with AMPK inhibitor Compound C improved the sensitivity of M059J glioma cells to irradiation, especially in relatively high dose (Figure 5B). It was suggested that acetylation of histone H3K14, H4K12 and H4K16 were important for Brd4 binding [59]. In consistency, JQ-1 reduced acetylation of H3K14 and H3K18 in both wildtype and AMPKα-KO cells. However, quite different effect of JQ-1 on the acetylation of H3K23 and H2BK12 was observed in wildtype and AMPKα-KO cells. JQ-1 treatment had no effect on H3K23 acetylation in wildtype cells but reduced H3K23 acetylation in AMPKα-KO cells. In contrast, JQ-1 treatment didn’t change H2BK12 acetylation in AMPKα-KO cells but reduced H2BK12 acetylation in wildtype cells (Figure 5C and D). Notably, JQ-1 reduced histone acetylation before irradiation, with no obvious changes observed after irradiation, suggesting JQ-1 regulated histone acetylation in irradiation-independent manner. Since wildtype and AMPKα-KO cells showed opposite trend on H2BK12 and H3K23 acetylation after JQ-1 treatment, we hypothesized AMPK signaling and BRD4 signaling coordinately regulated H3K23 and H2BK12 acetylation. Next, we investigated how the JQ-1-induced basal changes in histone acetylation finally resulted in lower cell viability under irradiation. Herein, we investigated whether repair signaling activated normally. JQ-1 treatment resulted in an enhanced phosphor-KAP-1(Ser824) signals and a prolonged γH2A.X signals sustained in AMPKα-KO cells, but without changes in phosphor-p53(Ser15) (Figure 5E). Therefore, we hypothesized that JQ-1 treatment affected the chromatin relaxation by disrupting the orchestrated regulation of histone acetylation during DDR. Accumulated unrepair DNA double strand breaks, as indicated by enhanced γH2A.X signals, would induce apoptosis. Consistently, we observed that JQ-1 treatment induced a higher proportion of apoptosis cells in the AMPKα-KO cells than in wildtype cells (Figure 5F). Thus, we concluded that knockdown of AMPK disrupted the orchestrated histone acetylation, which promoted the pro-apoptotic effect induced by JQ-1 treatment during DDR. Taken together, inhibition of AMPK activity improved the anti-tumor efficacy of JQ-1 via disturbing the balance in histone acetylation.

**Figure 5.**
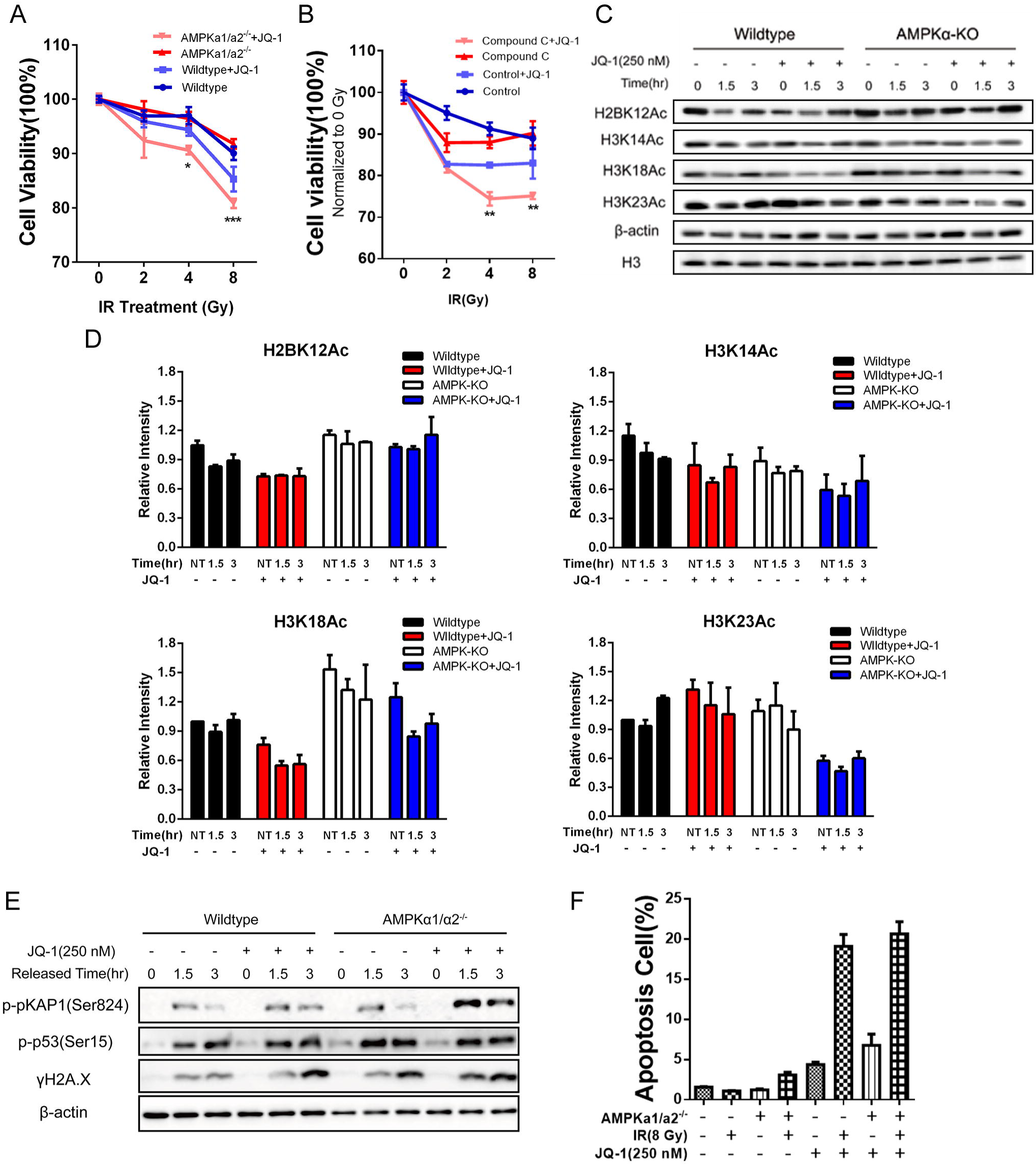
JQ-1 synergizing with AMPK inhibitor induces cell apoptosis in irradiation. **A**. JQ-1 treatment sensitizes AMPKα1/α2-deficient cells to irradiation. Both wildtype and AMPKα-KO cells were pretreated with 250 nM JQ-1 for 12 hours and then exposed to increasing dose of X-Ray (0, 2, 4, 8 Gy). After 72 hours the cell viability was detected by MTS assay. Each group was normalized to NT (0 Gy), respectively. **B**. JQ-1 synergizing with AMPK inhibitor sensitizes cells to irradiation. M059J cells were pretreated with 250 nM JQ-1 for 12 hours. 2 hours before irradiation, 1 μM Compound C was added to the culture medium. Then the cells were exposed to increasing dose of X-Ray (0, 2, 4, 8 Gy). After 72 hours the cell viability was detected by MTS assay. Each group was normalized to NT (0 Gy), respectively. **C**. AMPK knockdown disturbs the response of histone acetylation to JQ-1 treatment. Both wildtype and AMPKα-KO cells were pretreated with 250 nM JQ-1 for 12 hours and then exposed to a single dose of 10 Gy X-Ray. The cells were harvested at the indicated timepoint and analyzed by immunoblot. **D**. Quantitative result of panel C as measured by ImageJ software. Mean values ±SEM. from two independent experiments are shown. **E**. JQ-1 treatment prolonged the heterochromatin relaxation signals in AMPKα-KO cells. The samples collected in the same conditions as panel C. **F**. JQ-1 treatment synergizing with irradiation are prone to induce apoptosis in AMPKα-KO cells. Wildtype and AMPKα-KO cells were pretreated with 250 nM JQ-1 for 12 hours and then exposed to a single dose of 10 Gy X-Ray. 24 hours post irradiation cells were stained by PI and FITC-AnnexinV dyes and its mean fluorescence intensity was recorded by FACS. Mean values ±SEM. from two independent experiments. Data are normalized to the negative control (Wildtype without IR and JQ-1 treatment).

## Discussion

AMPK is an essential eukaryotic energy sensor to coordinate metabolism and other biologic activities. LKB1, one of the upstream kinases of AMPK, was a well-defined tumor suppressor [60]. However, fully understanding the exact role of AMPK seemed challenging. Previous works suggested AMPK, unlike its upstream kinase, played contradictory roles in tumor progression and therapy. AMPK activation regulated diverse downstream signaling pathways, together with complicated tumor microenvironment, which led to context-dependent cell fates. In one hand, AMPK phosphorylated ULK1 and promoted cell autophagy, to maintain cell survival in conditions of hypoxia and low nutrition [16, 61]. On the other hand, AMPK negatively regulated TSC-mTOR axis and inhibited cell outgrowth under the circumstance of energy deprivation [62-64]. It seemed that cell fate resulted from AMPK activation mainly depended on comprehensive signaling network within heterogenic tumor cells. The intercellular signaling network in types of tumors was balanced and maintained by both extrinsic stimulators like hypoxia and growth factors, as well as intrinsic features like gene signatures and transcriptional regulation [65, 66]. Genome instability and mutation was defined as one of the hallmarks in various types of cancer [67]. Abnormal DNA repair, especially dysfunctional DNA double strand breaks repair, impacts on the genetic fidelity and chromatin structure and consequently leads to genome instability and mutation [32, 68]. Previous works suggested AMPK was involved in DNA-damaging agent induced cell apoptosis but the underlying mechanism remained to be further understood [34, 69, 70]. Meanwhile, several studies shed light on the linkage between AMPK and DDR. DDR is a critical process to activate accurate repair pathways and other biological processes to coordinately repair damages. Therefore, a comprehensive understanding of the AMPK’s role in DDR is a critical step toward understanding the exact role of AMPK in tumor biology.

To address the question, we designed study to investigate whether novel AMPK substrates were involved in DDR. We carried out a system-wide phosphoproteomic study and global histone PTM analysis. Bioinformatic analysis of significant altered phosphoproteins enriched DNA damage repair-associated events and chromatin-associated functions, suggesting complicated regulatory network of AMPK in DDR. After comparing these altered phosphosites to classic AMPK substrates, we totally obtained 42 potential substrates of AMPK from basal group and irradiation group. This result suggested although irradiation might dramatically induce phosphorylation changes, AMPK likely had limited direct impact on DNA-associated signaling pathways.

Corresponding to the fact that chromatin-associated functions were enriched in phosphoproteomic study, histone modification analysis also suggested AMPK negatively regulated global histone acetylation. Therefore, an integrative analysis of two studies implicated a crosstalk between AMPK-mediated phosphor-regulation and histone acetylation. Based on this result, we further identified HDAC9, a member of Class IIa histone deacetylase family, as a new substrate phosphorylated by AMPK in an irradiation-independent manner. Overexpression of wildtype HDAC9 in cells increased the acetylation level at H2BK12 and H3K18, and the non-phosphorylable mutant HDAC9-S239A further promoted the increase of H2BK12 acetylation. Thus, we concluded that AMPK-mediated phosphorylation on HDAC9 might be required in balancing the histone acetylation at H2BK12, but the mechanism remained to be further investigated.

It was worth noting that AMPK-mediated phosphorylation at Ser479 of ASPP2 was required in p53-induced apoptosis in irradiation-dependent way. p53 plays distinct roles via different co-factors in DDR, including induction of cell cycle arrest or apoptosis. ASPP2 is the co-factor of p53 required in spontaneous induction of apoptosis and cooperates with p53 to suppress tumor growth [71, 72]. Previous studies suggested high level of ASPP2 sensitized cells to irradiation and DNA-damaging agents [73, 74]. Overexpression of non-phosphorylable mutant ASPP2-S479A moderately impaired apoptosis, suggesting that AMPK promoted apoptosis during DDR by phosphorylating ASPP2 at Ser479. The accumulation of unrepaired DNA damages activated apoptosis signaling pathway, thus avoiding hereditary of abnormal genetic information in mitosis. Taken together, AMPK-ASPP2 axis promoted apoptosis to prevent cells from genomic instability caused by sustained damages during DDR. Further study might focus on whether the phosphorylation site influenced protein interaction of ASPP2 complex and its correlation with cancer incidence or drug resistance.

Finally, our findings might reveal a crosstalk between AMPK activity and histone acetylation in tumor biology. Indeed, disturbing the histone acetylation by Brd4 inhibitor JQ-1 enhanced the sensitivity of cells to irradiation via induction of apoptosis. It was worth noting that except for H3K14Ac, JQ-1 also influenced acetylation at H2BK12, H3K23 and H3K18. Additionally, AMPKα1/α2-deficient cells were more sensitive than wildtype cells to JQ-1 treatment. These results suggested the potential role of AMPK in cancer therapy. First, AMPK activity might be using as a biomarker to predict the therapeutic response to acetyltransferase inhibitors or deacetylase inhibitors. Second, selective AMPK inhibitor might be useful to improve the sensitivity of tumor cells to radio-therapy or target-therapy.

Therefore, our study provided a source of AMPK-associated phosphorylation network and histone acetylation events, which might be helpful to understand the role of AMPK in DDR.

## Materials and methods

### Antibody and compound

The primary antibodies Phospho-Histone H2A.X (Ser139) (Catalog No. 2577), Phospho-p53 (Ser15) (Catalog No. 9284), Phospho-AMPKα (Thr172) (Catalog No. 2535), Phospho-Acetyl-CoA Carboxylase (Ser79) (Catalog No. 3661), Acetyl-CoA Carboxylase (Catalog No. 3662), AMPKα (Catalog No. 2532), AMPKα2 (Catalog No. 2757), AMPKβ1 (Catalog No. 12063), AMPKβ2 (Catalog No. 4148), GAPDH (Catalog No. 2118) and Myc-Tag (9B11) (Catalog No. 2276) were bought from Cell Signaling Technology (Danvers, MA, USA). The primary antibodies Phospho-KAP1 (Ser824) (Catalog No. ab70369), H2B-Acetyl-K12 (Catalog No. ab61228), H4-Acetyl-K16 (Catalog No. ab109463), H3-Acetyl-K14 (Catalog No. ab82501), H3-Acetyl-K18 (Catalog No. ab1191), H3-Acetyl-K23 (Catalog No. ab61234), 53BP2 (Catalog No. ab236448) were bought from Abcam (Cambridge, UK). The primary antibody HDAC9 (Catalog No. MA5-32820) was bought from Thermo Fisher Scientific (Waltham, MA, USA). The DYKDDDDK Tag (Catalog No. 018-22783) was bought from Wako (Osaka, JPN). The antibody β-actin (Catalog No. AM1021B) was bought from Abgent (San Diego, CA, USA).

Secondary antibody Alexa Fluor 555 labelled donkey anti-rabbit IgG (Catalog No. A31572) and Hoechst 33342 Solution (Catalog No. 62249) was bought from Thermo Fisher Scientific (Waltham, MA, USA). The secondary peroxidase AffiniPure Goat Anti-Mouse IgG (H+L) (Catalog No. 115-035-003) and Peroxidase AffiniPure Rabbit Anti-Goat IgG (H+L) (Catalog No. 305-035-003) were bought from Jackson ImmunoResearch Laboratories (West Grove, PA, USA). Chemiluminescent detection was completed with enhanced chemiluminescent (ECL) western blotting reagents (Catalog No. RPN2236, GE Healthcare, Chicago, IL, USA).

The compound JQ-1 (Catalog No. S7110) was bought from Selleck (Houston, TX, USA). The AMPK activator AICAR (Catalog No. A9978) and AMPK inhibitor (Catalog No. 171260) were bought from Sigma (MO, USA) and Millipore (Burlington, MA, USA), respectively.

### Cell culture and stable isotope labeling by amino acids in cell culture (SILAC) labeling

Wildtype MEFs and AMPKα1/α2-KO MEFs cell lines were cultured in DMEM with light lysine (^12^C_6_^14^N_2_-Lys) and arginine (^12^C_6_^14^N_4_-Arg), or heavy lysine (^13^C_6_^14^N_2_-Lys) and arginine (^13^C_6_^15^N_4_-Arg), respectively. The proteome labeling efficiency of heavy isotopic amino acids was > 98%, as determined by MS analysis. The HeLa and 293T cells were cultured in DMEM growth medium (Catalog No. 12100061, Gibco, Carlsbad, CA, USA) supplemented with 10% FBS (Catalog No. 10099, Gibco, Carlsbad, CA, USA). The M059J cells were grown in a medium containing a 1:1 mixture of DMEM and F12 medium (Catalog No. 11330057, Gibco, Carlsbad, CA, USA) supplemented with 10% FBS. All cells were cultured at 37°C with 5% CO_2_.

### Protein lysate preparation and in-solution digestion

Cells were harvested and washed with pre-cold phosphate-buffered saline (PBS). Then, cells were lysed with lysis buffer (8M Urea in 100 mM NH_4_HCO_3_) and subjected to sonication. After sonication, the lysates were clarified by centrifugation at 21,130 g for 10 minutes. Equal amounts of WT and AMPKα1/α2-KO MEFs cell lysate were mixed. The cell lysate mixture was reduced by 5 mM dithiothreitol (Catalog No. D0632, Sigma-Aldrich, St. Louis, MO, USA) at 56°C for 30 minutes. Then 15 mM iodoacetamide (Catalog No. I6125, Sigma-Aldrich, St. Louis, MO, USA) was added to alkylate the sulfhydryl groups. The extra iodoacetamide was eliminated using 30 mM cysteine. The protein extract was digested with trypsin (Catalog No. V5280, Promega, Madison, WI, USA) (trypsin: protein = 1:50) at 37°C for 12 hours. For complete digesting, additional trypsin (trypsin: protein = 1:100) was added for another 4 hours. Same amount of protein extract was digested with chymotrypsin (Catalog No. 11418467001, Roche, Basel, CHE) (chymotrypsin: protein = 1:50) at 25°C for 18 hours. Both tryptic and chymotryptic peptides were desalted through Waters SepPak C18 cartridges (Catalog No. WAT054960, Waters, Milford, MA, USA), vacuum-dried and stored at −80°C for further analysis.

### Histone extraction and in-gel digestion

Histone extraction was carried out as previously published [75, 76]. The isolated cells were lysed with extraction buffer (10 mM HEPES, 10 mM KCl, 1.5 mM MgCl_2_, 0.5% NP-40, 1× protease inhibitor mixture). Lysate was centrifuged at 1000 g at 4°C. Then, the pellets were washed and resuspended in 0.2 M H_2_SO_4_ overnight at 4 °C. The mixture was clarified by centrifugation, and the supernatant was collected for trichloroacetic acid precipitation. The precipitate was washed with pre-cold acetone for several times. The precipitate was dried completely at room temperature, and then dissolved in water for SDS-PAGE separation. Bands of histones (H1, H2A, H2B, H3, and H4) were excised and subjected to in-gel digestion. The tryptic peptides were analyzed by LC-MS/MS. After MS analysis, histone modifications of H2A, H2B, H3 and H4 were quantified by examined the peak area of modified peptide ions.

### Phosphopeptide enrichment

Phosphopeptide enrichment was carried out as previously published [51]. In brief, tryptic peptides were dissolved in loading buffer (6% TFA, 80% ACN, 1 M lactic acid), and then incubated with titanium dioxide beads (GL Sciences, JPN) at room temperature at a proportion of peptide: TiO_2_ = 4:1. The titanium dioxide beads were then washed with loading buffer for three times, wash buffer A (0.5% TFA, 30% ACN) for one time and wash buffer B (0.5% TFA, 80% ACN) for two times. The phosphopeptides were eluted from the beads with 15% NH_3_H_2_O and separated into six fractions.

### Nano-HPLC-MS/MS analysis

The peptides were analyzed by an EASY-nLC 1000 system (Thermo Fisher Scientific, Waltham, MA, USA) connected to an Orbitrap Q-Exactive mass spectrometer (Thermo Fisher Scientific, Waltham, MA, USA). Peptides were eluted from a reverse-phase C18 column (75μm ID, 3μm particle size, Dikma Technologies Inc., CA, USA) with a 70 minutes gradient of 7% to 80% buffer B (90% acetonitrile, 10% H_2_O, 0.1% formic acid) at a flow rate of 300 nL/min. Full MS spectra with an m/z range of 350–1300 were acquired with a resolution of 70,000 at m/z = 200 in profile mode. The AGC targets were 1.0e6 for full scan and 1.0e6 for MS/MS scan, respectively. Fragmentation of the 16 most intense precursor ions occurred at the HCD collision cell with a normalized collision energy of 28%, and tandem MS were obtained with a resolution of 17,500 at m/z = 200. Dynamic exclusion duration was set as 60 seconds.

### MS data analysis

MS/MS data were processed using MaxQuant software (version 1.5.3.2). Mus musculus database from Uniprot (release 2018_10_13, 53781 entries) with a reversed decoy database was used for data processing. For database searching, trypsin was set as the specific enzyme and the maximum number of missed cleavages was fixed at 2. Carbamidomethylation of cysteine residues was set as a fixed modification; oxidation of methionine and protein N-terminal acetylation were set as variable modifications. Phosphorylation of serine, threonine and tyrosine was set as variable modification for phosphosite analysis. Precursor mass tolerance for MaxQuant analysis was set to 4.5 ppm and MS/MS tolerance was set to 20 ppm. FDR thresholds for protein, peptide, and modification sites were all set as 1%.

### Bioinformatic analysis

DAVID bioinformatics functional annotation tool was used to identify enriched GO and KEGG pathway terms. Mus musculus genome was used as background in DAVID functional annotation analysis. The significance of fold enrichment was calculated using p-value < 0.05. Gene sets with p-value < 0.05 were visualized in enrichment maps using EnrichmentMapApp and Cytoscape. Motif-x was used to identify phosphorylation motifs present in significantly changed phosphoproteins.

### In vitro kinase assay and autoradiography

Briefly, the recombinant AMPKα2β2γ2 complex (200 nM) was pre-incubated with 20 nM CAMKKβ to be fully activated in reaction buffer (5 mM MgCl_2_, 20 mM Tris-HCl, 8 nM ATP, 1 mM DTT) at 37C for 1 hour. Then 200 nM substrate was incubated with 20 nM activated AMPK in reaction buffer at 37C for 1 hour, in which ATP was replaced by [^32^P]-labelled ATP (Catalog No. BLU002250UC, Perkin Elmer, Waltham, MA, USA). Afterwards, the reaction mixture was terminated by SDS loading buffer and subjected to immunoblot analysis Then SDS-PAGE was sealed with photographic film together for 12 hours before the film was fixed.

### Co-immunoprecipitation

Interested plasmids were introduced into 293 cells for 48 hours, then 293T cells over-expressing targeted proteins were lysed by buffer A (Catalog No. P0013B, Beyotime Biotechnology, Shanghai, CHN) (supplemented with 10 mM NaF, 1 mM Na_2_VO_3_ and protease inhibitor cocktails). The lysates were centrifuged at 10000 rpm, 4C for 10 minutes. The separated supernatant was divided into three fractions and respectively incubated with negative IgG or reciprocal antibodies at 4C overnights. Then pre-processed protein A agarose (Catalog No. P3476, Sigma, MO, USA) was added to mixture. After another 2-hours incubation, the beads-antibody-proteins complex was isolated from mixture by centrifuging at 1000 rpm, 4C for 5 minutes. To remove the non-specific binding proteins, the beads-complex was washed with pre-cold PBS buffer for three times before the samples were subjected to immunoblot assay.

### Cell viability assay

Both wildtype and AMPKα-KO MEFs cells (5000 per well) were pre-treated with 250 nM JQ-1 for 12 hours, followed by exposure to increasing dose of X-ray (0, 2, 4, 8 Gy).

M059J (7500 per well) were pre-treated with 250 nM JQ-1 and 1 μM Compound C for 12 hours, followed by exposure to increasing dose of X-ray (0, 2, 4, 8 Gy). After 72-hour recovery, the cells were subjected to MTS assay according to the manufacturer instructions. 10 μL per well of MTS/PMS (20:1, Promega, Madison, WI, USA) solution was added to each well containing 100 μL of culture medium, followed by a gentle shake. After incubation at 37 °C under 5% CO_2_ for 4 hours, the absorbance of the solutions was measured at 490 nm, using an M5 microplate reader (Molecular Device, San Jose, CA, USA).

### Cell Cycle Assay

Both wildtype and AMPKα-KO MEFs cells (7.5*10^4^/well) were exposed to a single dose of 8 Gy X-ray. After 24-hour recovery, cells were sampled by EDTA-free trypsin and 75% ethanol. Fixed cells were then incubated with 50 μg/mL propidium iodide (Catalog No. P4170, Sigma, MO, USA) and 100 μg/mL RNaseA (Catalog No. ST576, Beyotime Biotechnology, Shanghai, CHN) for 15 minutes at room temperature. The mean fluorescence intensity of DNA content was recorded by Flow cytometer (NovoCyte D2060R, San Diego, CA, USA) by PE channel. 10,000 events per sample were collected and analyzed by Software NoveExpress.

### Cell Apoptosis Assay

The cell apoptosis assay was performed by Annexin V-FITC/PI apoptosis detection kit (Catalog No. KGA108, KeyGEN BioTECH, Nanjing, CHN) according to the manufacturer instructions. Briefly, 2*10^5^ cells were prepared according to experimental design. At the sample point, cells were digested with EDTA-free trypsin and incubated with staining solution (500 μL Detection Buffer supplemented with 5 μL PI and 5 μL Annexin-V) for 15 minutes at room temperature. The fluorescence intensity were recorded by Flow cytometer (NovoCyte D2060R,, San Diego, CA, USA) by FITC and PE channel after fluorescence compensation deduction. 15,000 events per sample were collected and analyzed. Total apoptosis cells were FITC^+^PI^−^ cells plus FITC^−^PI^+^ cells. Data are presented as a densitometric ratio change that normalized to the negative control.

## Data availability

All mass spectrometry raw data have been deposited to the iProX, a full member of the ProteomeXchange consortium (Dataset ID: IPX0001446000). Reviewer can access data through website: https://www.iprox.org/page/SSV024.html;url=1579003580788LG0U Password: mKda

## Authors’ contributions

Jia Li, Minjia Tan and Yi Zang conceived the project and provided supervision; Yuejing Jiang and Xiaoji Cong performed the research and analyzed the data; Shangwen Jiang and Ying Dong conducted some of the experiments; Lei Zhao helped with data analysis and visualization. Yuejing Jiang, Yi Zang, Xiaoji Cong and Minjia Tan wrote the manuscript.

## Competing interests

The authors declare no competing interests.

## Aacknowledgments

This study was supported by National Natural Science Foundation of China (No. 81872888, No.81821005, No. 81673489, No. 31871414), the Special Project on Precision Medicine under the National Key R&D Program (No. 2017YFC0906600), the Shanghai Science and Technology Development Funds (16430724100), Key New Drug Creation and Manufacturing Program of China (2018ZX09711002-004, 2018ZX09711002-007), and K. C. Wong Education Foundation.

## Supplementary material

### Supplementary Figures

**Figure S1**

Sequence investigation of AMPKα1/α2-KO cell lines 1#, 2# and 3#. Genomic DNA was exacted from all wildtype MEFs cells and three AMPKα1/α2 double knockout cell lines and then subjected to DNA sequencing.

**Figure S2**

Scatter plot of the correlation analysis of two replicate experiments in basal and irradiation groups.

**Figure S3**

Motif enrichment of significantly downregulated phosphosites in basal group to indicate AMPKα-KO specificity. Motif-X algorithm embedded in MoMo software (version 5.1.0) was used for motif enrichment. Significance threshold for enrichment was set to 0.000001. Enriched p-value was shown below the sequence motif.

**Figure S4**

ASPP2 interacts with AMPKβ1. 293T cells were co-transfected with plasmids Flag-ASPP2 and Myc-AMPKβ1 for 48 hours and subjected to immunoprecipitation with anti-Flag antibody. The immunoprecipitated samples were examined by immunoblot, using anti-Myc and anti-Flag antibodies.

**Figure S5**

HDAC9 interacts with AMPKα2. 293T cells were co-transfected with plasmids Flag-HDAC9 and Myc-AMPKα2 for 48 hours and subjected to immunoprecipitation with anti-Flag antibody. The immunoprecipitated samples were examined by immunoblot, using anti-Myc and anti-Flag antibodies.

### Supplementary Tables

**Supplementary Table1 Phosphoproteomics data**

**Supplementary Table2 Histone modification data**

## Notes

### Competing Interest Statement

The authors have declared no competing interest.

